# Differential gene expression analysis reveals novel genes and pathways in pediatric septic shock patients

**DOI:** 10.1101/611947

**Authors:** Akram Mohammed, Yan Cui, Valeria R. Mas, Rishikesan Kamaleswaran

**Affiliations:** University of Tennessee Health Science Center, Memphis, TN, USA

**Author notes:** Corresponding Author Rishikesan Kamaleswaran, Ph.D., 50 N. Dunlap St, 4th Floor 491R, Memphis, 38103, TN, USA, E.

## Abstract

Septic shock is a severe health condition caused by uncontrolled sepsis. Advancements in the high-throughput sequencing techniques have risen the number of potential genetic biomarkers under review. Multiple genetic markers and functional pathways play a part in the development and progression of pediatric septic shock. Fifty-four differentially expressed pediatric septic shock gene biomarkers were identified using gene expression data from 181 pediatric intensive care unit (PICU) patients within the first 24 hours of admission. The gene expression signatures discovered showed discriminatory power between pediatric septic shock survivors and nonsurvivors types. Using functional enrichment analysis of differentially expressed genes (DEGs), the known genes and pathways in septic shock were validated, and unexplored septic shock-related genes and functional groups were identified. Septic shock survivors were distinguished from septic shock non-survivors by differential expression of genes involved in the immune response, chemokine-mediated signaling, neutrophil chemotaxis, and chemokine activity. The identification of the septic shock gene biomarkers may facilitate in septic shock diagnosis, treatment, and prognosis.

## Introduction

Septic shock is a life-threatening organ dysfunction caused by imbalanced host response to infection ^1^. Multi-omics sequencing technologies have increased the number of genetic biomarkers ^2^. Single or combination biomarkers are increasingly being analyzed and tested in the context of genes, RNA, or proteins ^3–6^ Many strategies for uncovering biomarkers exist, such as mass-spectrometry-based, protein arrays and gene-expression profiling. Furthermore, it has been demonstrated that multiple genes and immune system-related pathways participate in the development of pediatric septic shock ^7^

High-throughput technologies have enabled analysis of the expression of a number of genes and determine the activity of these genes in different conditions ^8^. Statistical testing and machine learning methods have been developed to successfully utilize the omics data for biomarker discovery ^2,9–18^.

The purpose of this study is to identify differentially expressed pediatric septic shock biomarkers using gene expression data. To this end, gene expression data from 181 samples from PICU within the first 24 hours were analyzed using multiple statistical testing methods to identify gene biomarkers. The gene expression profiles discovered by this statistical approach may lead to new insights into genetic biomarkers for successful septic shock diagnosis ^19^. Using functional gene-set enrichment analysis, we validated the known septic shock-related genes, pathways and functional groups, and identified the unexplored septic shock-related genes, and functional groups. The discovery of the potential gene biomarkers may provide effective septic shock diagnosis, treatment, and prognosis.

## Results

### Identification of Differentially Expressed Upregulated and Down-regulated Genes

Based on the preset criteria of an adjusted p-value < 0.05, a total of 54 genes from 21,731 were shown to be differentially expressed between the Septic Shock Survivor and Non-survivor samples, including 47 genes that were up-regulated and 7 genes that were down-regulated. Sixteen DEGs with a fold change of at least 1.5 is shown in Table 1 (For the complete list, refer to Supplementary File 1).

**Table 1:**
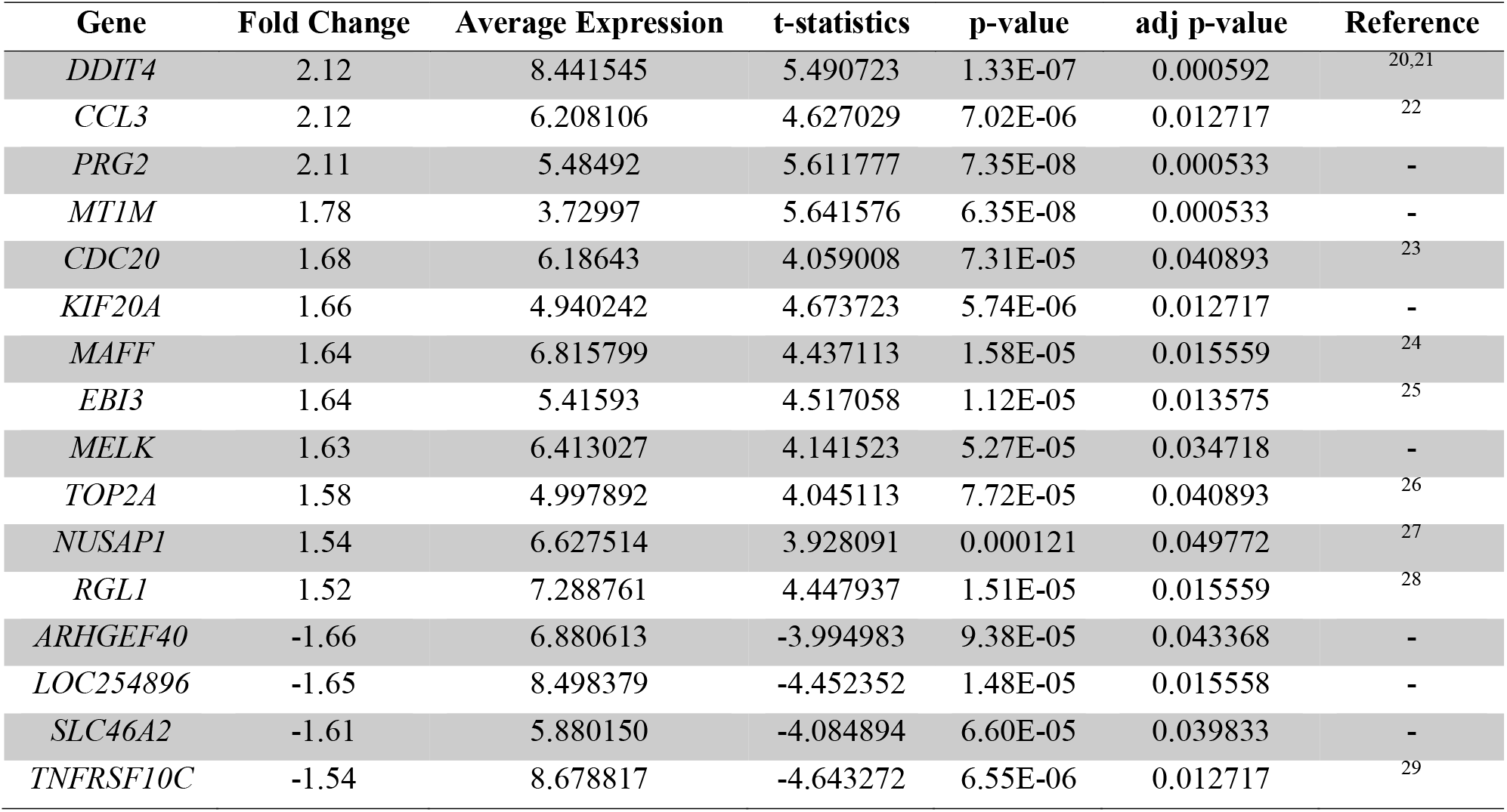
List of most significant up-regulated and down-regulated genes in septic shock

### Functional Enrichment Analysis of Differentially Expressed Genes

54 DEGs were analyzed by KEGG pathway and Gene Ontology (GO) term enrichment. A total of 52 genes were recognized in the DAVID database. KEGG pathway analysis revealed rheumatoid arthritis (RA) (hsa: 05323) and cell cycle (has: 04110) pathways as the most significant pathways (Table 2). GO analyses of the DEGs demonstrated that mitotic sister chromatid segregation (GO: 0000070), immune response (GO: 0006955), cell division (GO: 0051301), and chemokine-mediated signaling pathway (GO: 0070098) were the most enriched biological process (BP) terms (Table 2). ‘Chemokine activity (GO: 0008009) was the most enriched term under molecular function (Table 2). Chemokine interleukin-8-like domain (IPR001811), CC chemokine, conserved site (IPR000827) InterPro protein functional groups were among the significantly enriched functional classes associated with septic shock development (Table 2).

**Table 2:**
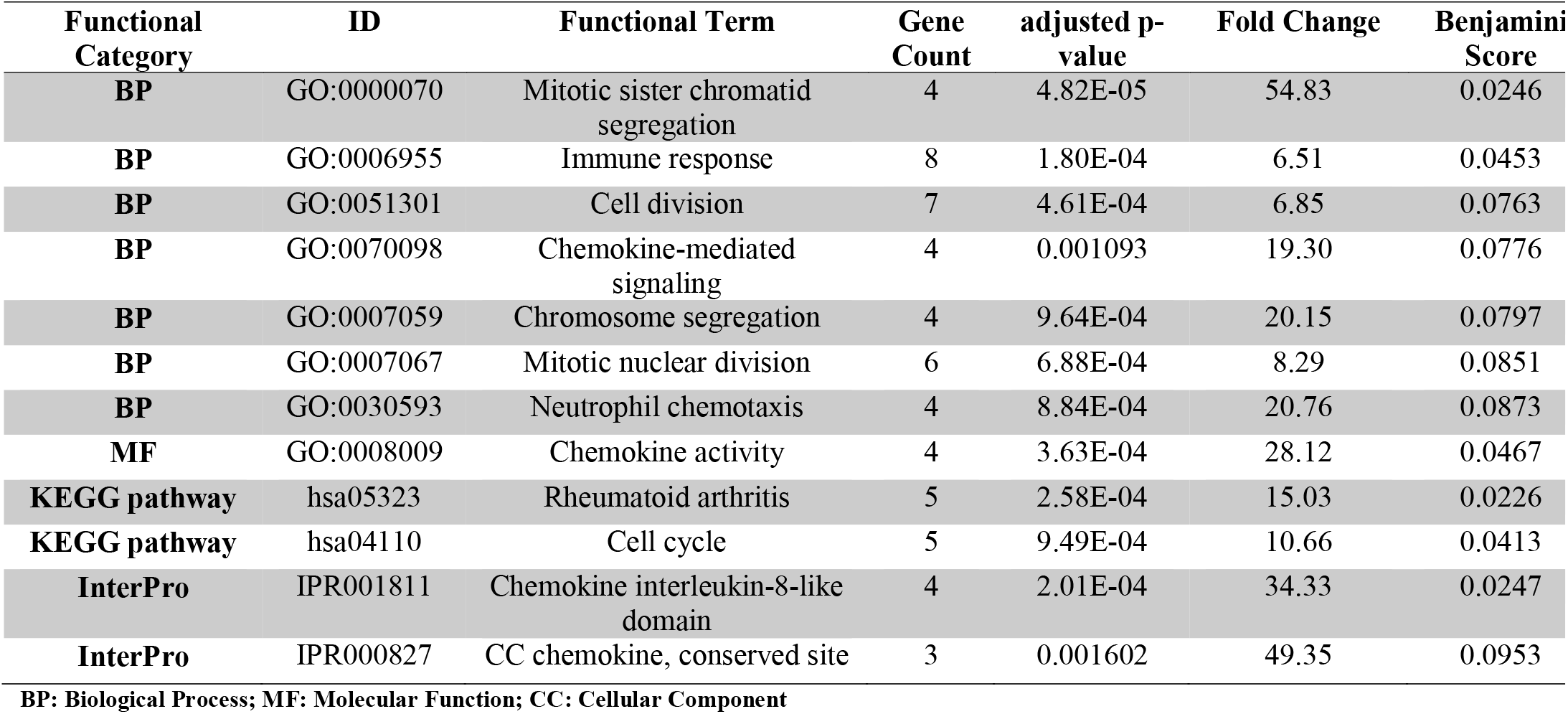
Functional enrichment of differentially expressed genes

## Discussion

This study of peripheral blood mRNA sequences revealed key genes and functional characterization associated with septic shock survivor and septic shock non-survival^30^. From the differential gene expression analysis, we identified the potential septic shock biomarkers that may help in an unbiased sepsis diagnosis, effective treatment, and ultimately improving prognoses. DEGs analysis using septic shock samples provides insights into the functional characterization of the genes between groups of septic shock survivor and non-survivor samples. However, like any other microarray data analysis is incomplete without performing adjustment for multiple testing. Due to approximately 20,000 (the approximate number of genes on a standard microarray chip) independent tests, it is expected to get at least 20 test scores by random chance when we allow a stricter p-value threshold of say 0.001. To avoid this situation, adjustment for multiple testing was utilized, and we have used the Benjamini Hochberg method. For a false-discovery rate (FDR) controlling procedure, the adjusted p-value of an individual hypothesis is the minimum value of FDR for which the hypothesis is first included in the set of rejected hypotheses, and we used an adjusted p-value cut-off of 0.05^31^.

We identified *CDC20* as one of the top up-regulated genes, along with *LCN2*, and *CD24*, similar to the findings of Dong et al., 2018 ^23^, which studied the development of trauma-induced sepsis in patients. However, our study population was much more diverse (Table 1). The most significantly up-regulated gene identified was *DDIT4* (DNA damage-inducible transcript 4-like). PERSEVERE-XP study had also identified *DDIT4* gene directly related to *TP53* ^7^. *DDIT4* (*REDD1*) is increased in the septic shock and can negatively regulate mTORC1 activity and plays an important role in energy homeostasis ^21^. We found *CCL3* as the second-most significantly up-regulated chemokine, a fundamental component of the acute-phase response to endotoxin in humans and regulates the leukocyte activation and trafficking ^32^. Elevated levels of *CCL3* has been detected within the first 24 hours of sepsis, suggesting its unique role in innate immune function ^22,33^. Further studies are needed to understand the mechanisms of these identified genes in the septic shock development. On the other hand, *TNFRSF10C*, a down-regulated gene has been shown to play an essential role in the sepsis immune response ^34^.

The set of genes identified is then examined for over-representation of specific functions or pathways. Septic shock survivors were distinguished from septic shock non-survivor by differential expression of genes involved in the immune response, chemokine-mediated signaling, neutrophil chemotaxis, and chemokine activity. Sepsis impacts the immune responses by directly altering the life span, production, and function of effector cells responsible for homeostasis ^35^. We identified the immune response (BP GO:0006955) term from DAVID analysis, and there has also been evidence that understanding the immune response to sepsis provides opportunities to develop effective treatment strategies ^36^.

Chemokines play a critical role in the sepsis and septic shock development, and molecules that block chemokine and chemokine receptor activity may prove to be useful in the identification of sepsis. ^37^. Our differentially expressed genes mapped to chemokine-mediated signaling (GO:0070098), chemokine activity (GO:0008009) molecular function, chemokine interleukin-8 like domain (IPR001811) and chemokine conserved site (IPR000827).

Sepsis and Rheumatoid Arthritis (RA) have been known to be associated for over 50 years ^38^. RA is shown to be a risk factor in sepsis patients, and sepsis infection could trigger the RA ^39^. We identified RA KEGG pathway (hsa:05323) using our differentially expressed gene set to be statistically significant (Table 2).

The gene expression changes shown in our results are based on the peripheral blood cells and may not be extrapolated as occurring at the organ or tissue level ^30,40^. Therefore, extra care must be taken while generalizing host immune responses or chemokine activities in septic shock patients. Besides, variations in the gene expression profiles of survivors and non-survivors of septic shock patients could be due to other unexplored confounding factors (such as patients demographics) rather than sepsis-related biology ^41^. On the other hand, the blood-based biomarkers have the advantage to be minimally-invasive. Large cohorts replication studies and network analysis studies are needed to gain insights into the relationships between these biomarkers and the survival/non-survival of cohorts ^42^. To avoid the possibility of selection bias the analysis must be expanded to the other independent data sets.

This work can be expanded by experimentally validating the identified blood-based biomarkers, developing robust machine learning methods to build septic shock prediction model using different omics data from diversified patient cohorts.

## Materials and Methods

### Data collection

Expression microarray data was collected from the NCBI Gene Expression Omnibus repository ^43^. The dataset contains the gene expression profiles of the peripheral blood samples from 181 septic shock patients including 154 survivors and 27 non-survivors, who were admitted to the pediatric intensive care unit within the first 24 hours ^44^. The GEO accession number for the data used in the study is GSE66099. The data was collected from the Affymetrix Human Genome HG-U133_Plus_2 (GPL570 platform).

### Normalization and Background Correction

The R Affy module ^45^ was used to remove the technical variations and background noise. The Quantile Normalization Method ^46^ was used to normalize the data, and the background correction was performed using the Robust Multi-Average ^47^ parameter method^48^.

### Probe to Gene Mapping

Affymetrix probes were mapped to the genes using the information provided in the Affymetrix database (hgu133plus2.db). We used average expression values when multiple probes mapped to the same gene ^19^.

### Identification of Differentially Expressed Genes

Differentially Expressed Genes (up-regulated and down-regulated genes) were identified using R limma package with a Benjamini-Hochberg (BH) correction method and the adjusted p-value of < 0.05 was used.

### Functional Analysis

We used DAVID ^49^ for functional enrichment analysis of the DEGs from samples of septic shock survivor or non-survivor. The biological process (BP), cellular component (CC), and molecular function (MF) were identified from the Gene Ontology database. For the GO functional groups, KEGG pathways, and InterPro functional terms returned from DAVID functional analysis; we considered an adjusted p-value threshold of ≤ 0.05 and gene count of 3 or more from this study.

### Statistical analysis

R programming language ^50^ is used for downloading the Affymetrix data and gene mapping using R Affy, and Bioconductor package. A Fisher-exact test was performed for determining statistical significance among the gene ontology terms and functional classes. Benjamini Hochberg multiple test correction method was used for calculating the differentially expressed genes.

### Data availability

The R scripts other related files used for data preprocessing, normalization and differential gene expression analysis are available from https://github.com/akram-mohammed/septic_shock_degs. The datasets generated and analyzed during the study are available upon request.

## Acknowledgments

We would like to acknowledge Timothy E. Sweeney and Hector R. Wang for providing the data.

## Author contributions

RK and AM conceived and designed the study, developed the method and performed the analysis. YC, VRM contributed to the analysis. All authors wrote and proofread the manuscript.

